# Early life experience sets hard limits on motor learning as evidenced from artificial arm use

**DOI:** 10.1101/2021.01.26.428281

**Authors:** Roni O. Maimon-Mor, Hunter R. Schone, David Henderson Slater, A. Aldo Faisal, Tamar R. Makin

**Affiliations:** WIN Centre, Nuffield Department of Clinical Neuroscience, University of Oxford, Oxford, UK; Institute of Cognitive Neuroscience, University College London, London, UK; Laboratory of Brain & Cognition, NIMH, National Institutes of Health, Bethesda, MD, USA; Oxford Centre for Enablement, Nuffield Orthopaedic Centre, Oxford, UK; Departments of Bioengineering and of Computing, Imperial College London, London, UK

## Abstract

The study or artificial arms provides a unique opportunity to address long-standing questions on sensorimotor plasticity and development. Learning to use an artificial arm arguably depends on fundamental building blocks of body representation and would therefore be impacted by early-life experience. We tested artificial arm motor-control in two adult populations with upper-limb deficiency: congenital one-handers – who were born with a partial arm, and amputees – who lost their biological arm in adulthood. Brain plasticity research teaches us that the earlier we train to acquire new skills (or use a new technology) the better we benefit from this practice as adults. Instead, we found that although one-hander started using an artificial arm as toddlers, they produced increased error noise and directional errors when reaching to visual targets, relative to amputees who performed similarly to controls. However, the earlier a one-hander was fitted with an artificial arm the better their motor control was. We suggest that visuomotor integration, underlying the observed deficits, is highly dependent on either biological or artificial arm experience at a very young age. Subsequently, opportunities for sensorimotor plasticity become more limited.

## Introduction

We move our hands in space with such apparent ease, yet the underlying process involves complex computations, representations and integration of information across multiple systems and modalities (Scott, 2004; Wolpert, 1997). Learning to move our limbs precisely and accurately begins *in utero*, where embryos have been documented refining arm-to-mouth reaching movements (Zoia et al., 2007). The trajectory of optimising reaching across infancy (Berthier & Keen, 2006; Leed et al., 2019) and childhood (Contreras-Vidal, 2006; Schneiberg et al., 2002; Simon-Martinez et al., 2018; Sveistrup et al., 2008) is highly protracted, roughly plateauing at around 10-12 years of age. In the present study, we investigated reaching behaviour in two groups of individuals who experienced a vastly different motor development but share current motor constraints: individuals born with a partial upper-limb (missing a hand and a part of their arm; hereafter one-handers) and individuals who were born with a fully developed upper-limb but lost it as adults (hereafter amputees). We asked how a sensorimotor system that developed with (amputees) or without (one-handers) experience of a complete arm supports the control of an upper-limb substitute (artificial arm). Artificial arm motor control provides a unique opportunity to address key questions surrounding sensorimotor plasticity. The flexibility needed to support this new body part is arguably different from that observed in traditional motor learning paradigms (e.g., involving tools) as it might relate to more fundamental building blocks of body representation and the internal models for motor control.

We consider three possible predictions, involving differences in artificial arm motor control across these two groups: First, perhaps the most straightforward prediction is that one-hander’s artificial arm motor control would be superior to that of amputees. It is often thought that the brain is more plastic during earlier stages of development (Knudsen, 2004). Therefore, it becomes more difficult to acquire radically new motor skills in adulthood, which is probably why most virtuoso musicians and athletes started practicing their trade in their childhood (Penhune, 2011). As mentioned above, one-handers start using artificial arms at a very young age (in our sample as early as 3 months with an average of ∼2.5 years), even before early training for musical and athletic skills. It therefore stands to reason that in comparison to amputees, who only begin to learn to use their artificial arm as adults (in our sample at a mean age of 32), one-handers should have had more time and practice in early childhood to perfect their artificial arm motor skill. Moreover, amputees often experience a ‘phantom hand’ (Stankevicius et al., 2020), rooted in a maintained representation of their missing arm (Bruurmijn et al., 2017; Kikkert et al., 2016; Wesselink et al., 2019) which might in theory interfere with the acquisition of a representation of an arm substitute (the artificial arm). Perhaps most importantly, relative to amputees, one-handers tend to make better use of their artificial arm in daily life (Biddiss & Chau, 2007). Together, these considerations lead to a strong hypothesis that one-handers would have had better opportunities for developing sensorimotor artificial arm control.

A second alternative hypothesis is that one-handers’ early-life disability might offset motor development, but that such disability-related impairment would not necessarily lead to inferior motor performance with an artificial arm. It could be argued that regardless of the undisputed role of early life experience in shaping brain organisation and function, the canonical brain infrastructure will still exist and be able to support the dormant function, even in one-handers. In the visual domain, children born with high density cataracts who received corrective surgery later in life have been shown to retain some rudimentary forms of visual perception (Gandhi et al., 2017). This hypothesis is consistent with recent studies emphasising normal visuomotor processes and representations of hands of individuals born with no hands (Vannuscorps & Caramazza, 2015, 2016 though see Maimon-Mor, Schone, et al., 2020; Philip et al., 2015; Philip & Frey, 2011; Wesselink et al., 2019). Moreover, considering these individuals potentially have a lifetime of daily experience controlling an artificial arm, it is possible that they will be able to ‘close the gap’ that had started in early development, relative to their able-bodied peers. Indeed, it has been consistently shown how well adults can adapt their motor behaviour to overcome a myriad of perturbations, and learn to perform intricate and skilful tasks (Wolpert et al., 2011).

A third hypothesis asserts that experience with a complete arm early in life might be crucial for the successful integration of any arm, including an artificial one. Therefore, motor control of an artificial arm would be superior in acquired amputees who had ‘typical’ motor development for their missing arm, relative to one-handers who had atypical motor development. This idea is rooted in the old debate of the relative contributions of nature vs. nurture. Current views consider neural development an interaction between predetermined maturation based on a genetic template and experience (Adolph & Franchak, 2017; Karmiloff-Smith, 1998; Krubitzer & Prescott, 2018). The neural topographical organisation of sensory input across the cortex has been shown to be in part determined by genetics (Miyashita-Lin et al., 1999; Rubenstein et al., 1999). However, for both the motor system and the visual system (an integral input to the sensorimotor loop), early deprivation has been shown to have a permanent effect on development (de Heering et al., 2016; Walton et al., 1992). As such, individuals who, prior to their amputation, benefited from a typical developmental trajectory might be able to rely on the existing upper-limb infrastructure, after amputation, when learning to control an artificial arm. This is in stark contrast to one-handers, who never developed an upper limb, due to developmental malformation, and therefore lack both visual and motor experience of their missing limb during the formative years of their motor development. Based on this hypothesis, amputees would have superior motor control of artificial arms, compared to one-handers.

Consistent with this final hypothesis we found that although they had started training to use an artificial arm earlier in life, and sustained more elapsed years of artificial arm use, one-handers were unable to refine their reaching control to normal levels. One-handers produced larger reaching errors with their artificial arms compared to both artificial arm reaches of amputees and non-dominant hand reaches of age-matched controls. We used numerous measures and tasks to interrogate the potential contributors to sensorimotor performance across groups, allowing us to disentangle the different components that might have contributed the aforementioned group differences. Lastly, we explored how the key components contributing to reduced motor control relate to early life experience with an artificial arm. Our results suggest that the formation of an arm representation in early life has a long-lasting effect on the incorporation of an artificial arm, highlighting that opportunities for sensorimotor plasticity becomes more limited with age, even across early childhood.

## Results

### 1. One-handers show inferior artificial arm motor control

In order to assess motor control of artificial arms, participants performed visually guided reaches to a set of targets using a robotic manipulandum device (see Figure 1A&B). Motor control measures were compared across three groups: one-handers (n=18), amputees (n=14) and age-matched controls (n=19). All included participants were able to control the robotic handle and perform the task using the same speed-accuracy trade-off parameters following Fitts’ Law (see Supplementary Results). Reaching performance was evaluated by measuring the mean absolute error participants made across all targets (see Figure 1C). The absolute error refers to the distance from the cursor’s position at the end of the reach (endpoint) to the centre of the target in each trial. Participants completed the same task using their intact arm as well, allowing us to control for individual differences relating to aspects of the task that are not artificial arm specific. We found no significant group effect (*F*_*(2,48)*_=1.05, *p*=-.14) when comparing the absolute-errors of the intact arm and dominant arm across the three groups (controls, amputees and one-handers). However, this result was inconclusive (*BF*_*10*_=0.65), i.e., supports neither the null nor the alterative hypothesis. We therefore included the intact arm as a confound regressor in subsequent analyses (see result section 6 for additional results and discussion regarding intact arm performance).

**Figure 1.**
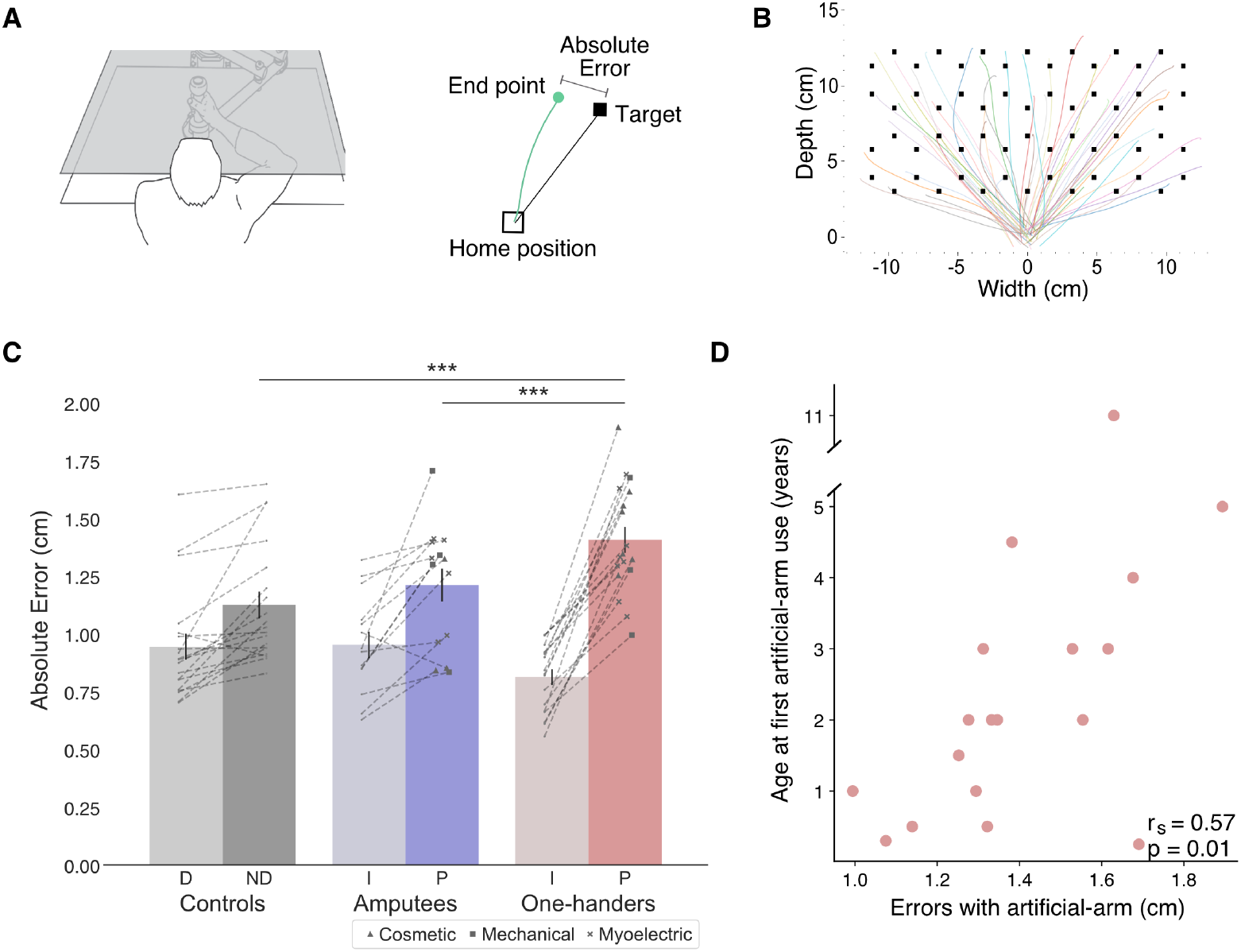
Experimental design and main analyses. (A) Left: An illustration of the robotic manipulandum device setup. Participants performed reaching movements while holding a robotic handle. A monitor displaying the task components was viewed via a mirror, such that participants did not have direct vision of their arm. Visual feedback was provided as a cursor depicting the current location of the arm. Right: A visualisation of a single trial and the different terms used. In each trial, participants reached from the home position to a single visual target. The green line represents the participant’s arm trajectory. (B) Reaching trajectories to all targets from a randomly selected participant. The different coloured lines are trajectories of individual reaching trials. (C) Reaching performance as measured by absolute errors for each group for each arm. Grey, blue and red colours represent controls, amputees and one-handers respectively. Lighter colours represent intact/dominant-arm performance; darker colours represent artificial/nondominant-arms. We found a significant group effect (*F*_*(2,47)*_=13.81, *p*<0.001, *η*_*p*^*2*^_=0.37), with one-handers making larger errors with their artificial arm compared to both amputees’ artificial arms (*t*=-3.77, *p*_*tukey*_=0.001, *Cohen’s-d* =-1.39), and controls’ non-dominant arm (*t*=-5.06, *p*_tukey_<0.001, *Cohen’s-d* =-1.705). Dotted lines connect errors between arms of individual participants. Artificial arm markers represent artificial arm type. (D) Relationship between age at first artificial arm use and artificial arm reaching errors in one-handers. Illustration in figure 1A was reproduced from Figure 1A, Wilson, Wong, & Gribble 2010, PLoS ONE, published under the Creative Commons Attribution 4.0 International Public License (CC BY 4.0; https://creativecommons.org/licenses/by/4.0/). D – Dominant arm, ND – Nondominant arm, I – Intact arm, A – Artificial arm. *** *p* < .001

We performed an analysis of covariance (ANCOVA) on participants’ artificial arm errors where participants’ intact arm errors were defined as a covariate, and group (controls, amputees, and one-handers) as a between-subjects variable. We found a significant effect of intact-hand performance (*F*_(1,47)_=28.65, *p*<0.001, η_p_^2^=0.38), i.e., participants who had small errors with their intact arm also tended to have smaller errors with their non-dominant/artificial arm. We found a significant group effect (*F*_(2,47)_=13.81, *p*<0.001, η_p_^2^=0.37), indicating the groups differed in their visuomotor performance with their artificial arm (or non-dominant arm in controls). Post-hoc comparisons revealed that one-handers exhibited larger errors with their artificial arm compared to both the artificial arm of amputees (*t*=-3.77, *p*_*tukey*_=0.001, *Cohen’s*-*d* =-1.39), and the non-dominant arm of controls (*t*=-5.06, *p*_*tukey*_<0.001, *Cohen’s*-*d* =-1.705). Conversely, amputees’ artificial-limb errors did not differ from those of controls’ non-dominant arm (*t*=-0.885, *p*_*tukey*_=0.65), indicating a specific deficit in error reaching for one-handers’ artificial arm. To further explore the non-significant performance difference between amputees and controls, we used a Bayesian approach (Rouder et al., 2009), inputting the smaller effect size of the two reported here (1.39) as the Cauchy prior width. The resulting Bayesian Factor (*BF*_*10*_=0.28) provided moderate support to the null hypothesis (i.e., smaller than 0.33). To summarise, one-handers show significantly inferior artificial arm reaching accuracy in our task compared to the other groups (see Supplementary Table S1 for results of the statistical analyses confirming that this effect was not driven by the side (L/R) of the artificial arm/non-dominant side).

### 2. Physical aspects of artificial arm use do not correlate with endpoint errors

We first wanted to rule out the influence of two crucial physical aspects of artificial-limb use: residual-limb length and device type. The length of the residual-limb, used to carry and control the artificial arm, can have a potential impact on the level of its motor control. The shorter the residual limb, the more restrictive the artificial-limb control is, e.g., due to more restrictive motion and less leverage. Across both amputee and one-hander groups, the correlation between absolute reaching error and either residual-limb length was not significant (*r*_*s*_(30)=-0.23, *p*=0.2). Moreover, repeating the previously reported ANCOVA analysis while adding residual limb length as a covariate revealed no significant effect of residual-limb length (*p*>0.2) and, importantly, did not abolish the group effect (*p*=0.03; for a full statistical report see Supplementary Table S2). Therefore, the length of the residual limb does not play a significant role in the observed group effect for end-point accuracy of artificial arm reaches.

It is important to note that while artificial arm devices have different levels of wrist- and grasp-control, they are all used similarly during reaching (i.e., in the current task the participants did not use any of the devices additional control features). Yet, we wished to confirm that differences in artificial arm types used across the two groups did not affect our findings. The devices used by our participants can generally be categorised into three device-types: (1) ‘cosmetic’ devices that look like a hand (*n*=10), these are static devices that do not afford additional control, (2) ‘body-powered’ devices in the shape of a hook (*n*=7), these include a mechanical grip control, (3) ‘myoelectric’ devices (*n*=15), these are relatively heavy devices, controlled using signals from the muscles of the residual-limb and powered by motors to perform grip functions (see marker type in Figure 1C). Despite the differences in appearance and control mechanisms between devices, the type of device used does not seem to influence endpoint reaching error in our task, as demonstrated by a one-way ANOVA with device type as a between subject variable showing no significant differences between devices (*F*_(2,29)_=0.435, *p*=0.65, *BF*_*10*_=0.275).

### 3. One-handers’ artificial arm errors originate from increased motor noise

In our analyses so far, we reported the absolute error – the average distance of the endpoint from the visual target across all reaches. An increase in absolute error can be the result of two different type of error components (see Figure 2A): bias (e.g., consistently reaching to the left of the target) and noise (variability/spread of endpoints). These are often also referred to as accuracy and precision, respectively. A larger bias is caused by a model-mismatch, for example, an inaccurate internal forward model that produces a biased control policy that consistently fails to accurately transport the arm to the correct location, resulting in poor accuracy. Several different sources can cause a noisier performance, for example: large uncertainties in the sensory estimates of proprioception (Gordon et al., 1995), motor noise (Faisal et al., 2008), or a result of a failed computation (Contreras-Vidal, 2006), e.g. to optimally use sensory inputs to reduce this inherent noise. Assessing these error components separately can give us an insight into the underlying processes that are affected in one-handers.

**Figure 2.**
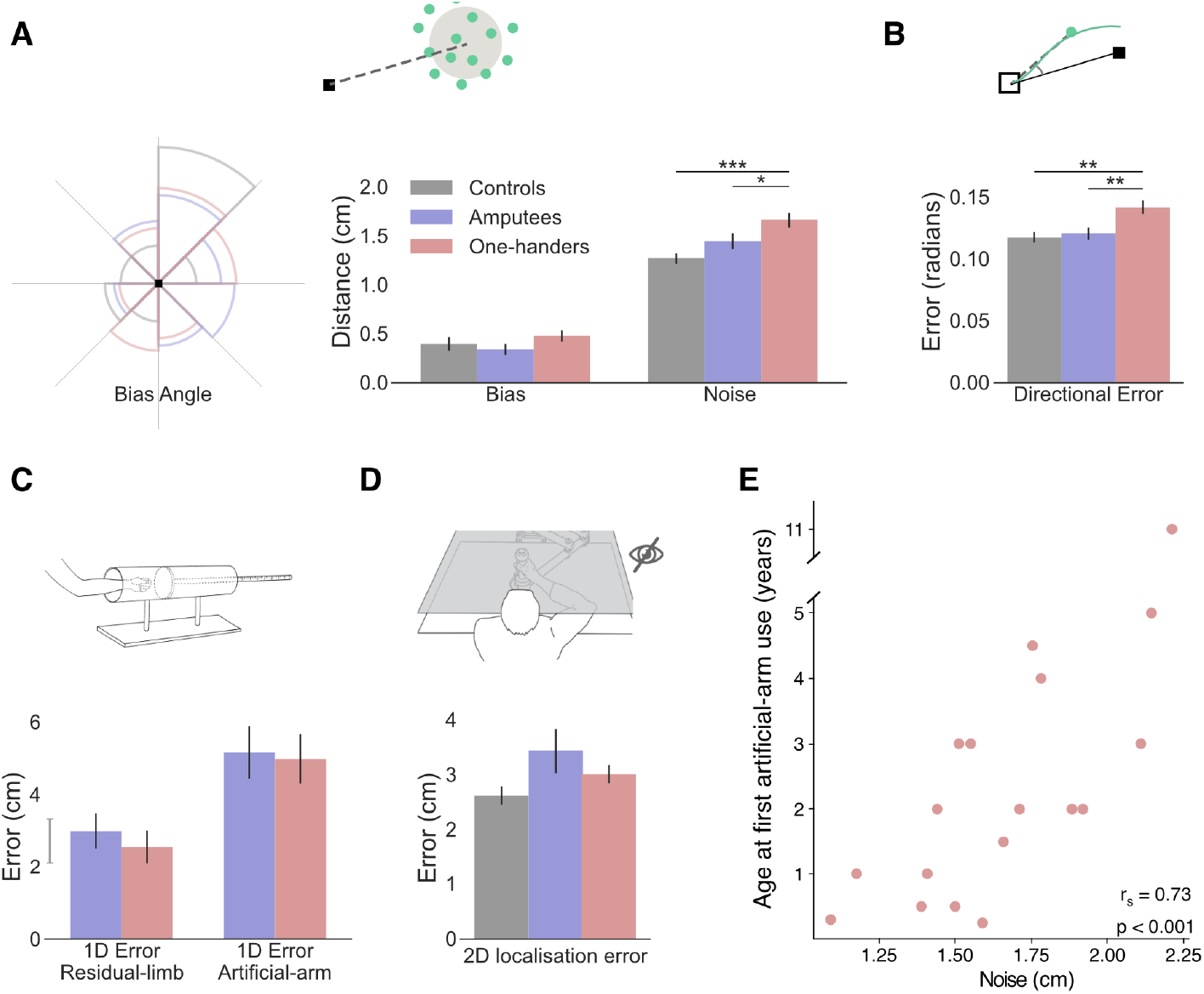
Exploring the source of increased reaching errors using additional analyses and tasks. In all plots, grey, blue and red represent controls, amputees and one-handers (respectively). (**A**) Left: rose plot density histogram of the distribution of bias angles across the groups, the larger the arc the more individuals from that groups had a bias within the arcs angle range. We found no significant differences in bias angle between the groups (Watson-Williams circular test: *F*_(2,48)_=1.95, *p*=0.15). Right: Error bias and noise results. No significant group differences were found for bias (*F*_(2,47)_*=*2.40, *p*=0.1, *BF*_*Incl*_=0.72). One-handers show significantly more motor noise than amputees and controls (*F*_(2,47)_=14.15, *p*<0.001, η_p_^2^=0.38; post-hoc significance levels are plotted). (**B**) Initial directional error results. One-handers have larger directional error in the initial phase of reaching (*F*_(2,47)_=8.01, *p*<0.001, η_p_^2^=0.26; post-hoc significance levels are plotted). (**C**) 1D localisation task results. Participants placed their residual-limb or artificial arm inside an opaque tube and were asked to assess the location of the limb using their intact arm. We found no localisation differences between amputees and one-handers in either condition (*BF*_*10*_<0.33 for both). The grey line next to the y-axis shows the mean ± s.e.m of controls non-dominant hand localisation errors. (**D**) 2D localisation task results. Using the same apparatus, participants performed reaches to visual targets without receiving visual feedback during the reach. We found no group differences in absolute error (*F*_*(2,44)*_=0.71, *p*=0.5, *BF*_*Incl*_=0.33). (**E**) Relationship between artificial arm motor noise and age at first artificial arm use artificial arm in one-handers. Illustration in figure 2D was reproduced from Figure 1A, Wilson, Wong, & Gribble 2010, PLoS ONE, published under the Creative Commons Attribution 4.0 International Public License (CC BY 4.0; https://creativecommons.org/licenses/by/4.0/).. *\* p* < .05, *\*\* p* < .01, *\*\*\* p* < .001

In order to calculate these measures across all targets, we drew the error vector for each trial (the line connecting the target with the end-point location) and overlaid all the error vectors, as if they were made to a single target. While error vectors are known to be location-dependent (Van Beers et al., 1999, 2004), because we compare bias and noise measures across groups, and the distribution of error vector directions did not differ between groups (Watson-Williams circular test: *F*_(2,48)_=1.95, *p*=0.15; see Figure 2A), these measures are suitable for present purposes. We compared artificial arm and nondominant arm biases (distance from the centre of the endpoint to the target) across groups, using intact arm biases as a covariate. The ANCOVA resulted in no significant group differences (*F*_(2,47)_*=*2.40, *p*=0.1, *BF*_*Incl*_=0.72; see Figure 2A). When comparing artificial arm and nondominant arm motor noise (spatial standard deviation (SD) of end-points relative to the centre of the endpoints), using intact-hand noise as a covariate, we found a significant group effect (*F*_(2,47)_=14.15, *p*<0.001, η_p_^2^=0.38; see Figure 2A). Reflecting the absolute error findings, one-handers exhibited larger motor noise with their artificial arm compared to the artificial arm of amputees (*t*=-2.90, *p*_*tukey*_=0.015, *Cohen’s*-*d*=-0.855), and the non-dominant arm of controls (*t*=-5.31, *p*_*tukey*_<0.001, *Cohen’s*-*d*=-1.65). Comparing motor noise between amputees’ artificial arm and controls’ non-dominant arm was inconclusive (*t*=-2.1, *p*_*tukey*_=0.1, *BF*_*10*_=1.2). Similar results were obtained when testing for the unique effect of noise beyond bias, by adding artificial arm bias as a covariate when comparing the motor noise of the artificial arm errors between the three groups (See Supplementary Table S3 for a full statistical report). These results show that one-handers’ artificial arm reaches are best characterised by increased noise (end-point variability). In the next analyses, we will test two potential sources of noise: artificial arm sense of localisation (proprioception) and adequacy of motor planning and its execution.

### 4. One-handers and amputees are equally accurate at localising their artificial arm without visual feedback

Commercially available artificial arms, as the ones used by our participants, currently lack direct sensory feedback, and of most relevance to reaching, proprioceptive feedback. Proprioception, the sense of position and movement of our body, provides an essential input to the sensorimotor system (Sarlegna & Sainburg, 2009; Wolpert et al., 1995). Together with vision it is used to accurately localise the current position of the arm and guides corrective movement during the execution of reaching movements. The lack of proprioception is therefore a reasonable candidate for explaining the inferior control of the artificial arm. As a proxy measure for proprioception, we assessed artificial arm localisation abilities. First, to assess artificial arm localisation, at its most basic and simple form, we tested artificial arm localisation along a single axis. In a separate task, participants were asked to place their artificial arm in an opaque tube and use their intact arm to point to the end-point of the artificial arm (McDonnell et al., 1989); see Methods). Since the artificial arm is sensed and localised via the residual-limb, we also assessed the proprioception of the residual-limb. Interestingly, we found no localisation differences between amputees and one-handers in either condition (Residual-limb: *Mann-Whitney W*=112.5, *p*=0.46, *BF*_*10*_=0.26; Artificial arm: *Mann-Whitney W*=104.5, *p*=0.7, *BF*_*10*_=0.24; See Figure 2C), suggesting that one-handers are equally able as amputees at localising their artificial arm.

While artificial arm localisation does not seem to differ between our two limbless groups, it is still possible that the online integration of localisation input, rather than the input itself, is suboptimal in one-handers. To test this, we asked participants to perform a task very similar to our main reaching task, with the exception that participants reached to visual targets without receiving continuous visual feedback of their limb position (see Methods). Here, participants were instructed to prioritise accuracy in their performance. Using the same ANCOVA approach described above, we compared artificial arm (and non-dominant arm) errors of the three groups, while controlling for intact-hand performance as a covariate. We found no group differences in artificial arm errors (F_(2,44)_=0.71, *p*=0.5, BF_Incl_=0.33; See Figure 2D). We further performed a planned pairwise comparison between artificial arm performance of one-handers and amputees and found no significant difference (*t*=0.71, p_unc_=0.48, *BF*_*10*_=0.25). Together, these results suggest that one-handers’ artificial arm localisation is not substantially different than that of amputees.

### 5. One-handers show larger artificial arm initial directional errors

Fast reaching movements, such as the ones performed in our main task, can be roughly divided into two phases: an *initial impulse phase* that involves the execution of a motor plan constructed prior to movement initiation and an *error correction phase* where sensory information is used to correct errors during execution (Elliott et al., 2001). The timing of peak velocity in such a movement is often used as a time-point that mostly reflects the first phase, i.e. the trajectory up to this point is mostly governed by feedforward mechanisms (Krakauer et al., 1999; Patterson et al., 2017; Sainburg et al., 2003). As the error at the end of the reach can originate from both feedforward processes and sensory integration processes, comparing the errors at the initial phase allows us to disentangle feedforward from feedback mechanisms. Specifically, the directional error at this stage provides a measure of how far away the movement is from the target’s direction. To measure the initial directional error for each trial we took the direction vector (the line connecting the home position and the arm’s location at peak velocity; see Figure 2B) and calculated the absolute angle between that direction vector and the target direction vector (the line connecting the target and the home position). While reaching paths are known to diverge from the straight line differently depending on the reach direction (Van Beers et al., 1999, 2004), because we compare directional errors across groups, this measure is suitable for our present purposes. For this reason, we also do not assume the ‘optimal’ directional error would be zero. We use the initial directional error to characterise the early part of the arm trajectory as an indicator of the accuracy of the motor plan. As with previous measures, initial directional errors were analysed using an ANCOVA comparing directional errors of the artificial arm (and non-dominant arm) of the three groups, while controlling for intact-hand directional errors as a covariate. We found significant group differences in artificial arm errors (*F*_(2,47)_=8.01, *p*<0.001, η_p^2^_=0.26). One-handers exhibited larger directional errors with their artificial arm compared to amputees (*t*=-3.515, *p*=0.003, Cohen’s-*d*=-1.10), and controls (*t*=-3.31, *p*=0.005, Cohen’s-*d*=-0.98). Amputees’ artificial arm directional errors did not differ from those of the nondominant arm of controls (*t*=.16, *p*=0.9, *BF*_*10*_=0.21). This result suggests that one-handers’ initial motor plan differs from that of amputees and controls. However, based on this measure alone, we are unable to distinguish between errors resulting from the motor plan itself or with noise resulting from its execution.

### 6. Early-life but not present experience with artificial arms effects current motor control in one-handers

We tested the hypothesis that present artificial arm usage will have a significant relationship on users’ artificial arm motor control. First, we confirmed that although one-handers have accumulated on average ∼29 more years of (intermittent) artificial arm experience compared to amputees (*t*_(30)_=-7.86, *p*<0.001), there are no differences in artificial arm daily usage between the two artificial arm users’ groups, as assessed using questionnaires (acquired amputees vs one-handers; *t*_(30)_=-0.25, p=0.81, BF_10_=0.35). Contrary to our hypothesis we found no such relationship between artificial arm reaching errors and a daily-life artificial arm usage score, encompassing both daily wear-time and functionality of use (*r*_(30)_=-0.05, *p*=0.78, *BF*_*10*_=0.23). We did, however, find a relationship between daily-life artificial arm usage and intact-hand reaching errors across amputees and one-handers (*r*_(39)_=-0.41, *p*=0.008; see Supplementary Figure S1). Smaller intact-hand errors (higher accuracy) were associated with higher artificial arm use scores (more versatile and frequent use). While being cautious not to infer causality from a correlation, we believe this result uncovers a relationship between an individual’s general motor control (as measured by their intact arm) and their ability to use an artificial arm. This result further highlights the need to control for individual differences in intact-hand motor control when studying artificial arms.

While we found that artificial arm present-day use does not predict its motor control (i.e., absolute error in reaching), we wanted to next test whether the user’s early-life experience does. Quantifying something as complex as artificial arm past use is a difficult feat. Here, we focused on the age at which one-handers were first fitted with an artificial arm (range: 2 months – 11 years). Interestingly, we found a significant positive correlation between age at first artificial arm use and artificial arm reaching errors (*r*_*s*(16)_=0.57, *p*=0.01; see Figure 1D). One-handers who started using an artificial arm at an earlier age produced smaller errors with their current artificial arm as adults. This result suggests that our ability to adjust our motor representation might not be as flexible as we thought and might be constrained by early life experience. Finally, we did not find a significant relationship between past (age at first artificial arm use) and present (daily-life use score) artificial arm experience (*r*_*s*(16)_=-0.03, *p*=0.91), or between artificial arm reaching errors and elapsed time since first artificial arm use (*r*_(16)_=0.01, *p*=0.96, *BF*_*10*_=0.29).

### 7. One-handers age at first experience with an artificial arm correlates with motor noise

Next, we wanted to test which of the aforementioned measures, each representing a different aspect of motor control, best correlates with age at first artificial arm use. Discovering which of these measures are influenced by age at first use can give us an insight into which age-constrained motor control process might be involved in learning to control an artificial-limb. Motor noise and initial directional error, being the two measures that produced a significant group effect, are of special interest, but for the sake of completeness we have tested the potential impact of all six measures. We found that only motor noise significantly correlated with age at first artificial arm use (*r*_*s(16)*_=0.73, *p*_*bonf*_=0.003 [corrected for 6 comparisons]; see Figure 2E). That is, individuals who started using an artificial arm earlier in life also showed less end-point motor noise in the reaching task. While we did find group differences in initial directional errors (as reported in result section 5), it did not significantly correlate with age at first artificial arm use (*r*_*s(16)*_=0.38, *p*_*bonf*_=0.74). From these results, we infer that early-life experience relates to a suboptimal ability to reduce the system’s inherent noise, and that this is possibly not related to the noise generated by the execution of the motor plan. Therefore, improved motor control, due to early life experience, might relate to suboptimal integration of visual and sensory information within the sensorimotor system.

### 8. Amputees’ artificial arm control might also be constrained by a time-sensitive process following amputation

Finally, we wanted to explore whether amputees also show a parallel phenomenon of an effect of early-life experience of artificial arm use on current motor control. From the relationship observed in one-handers, we can draw two predictions with regards to amputees: first, that the age at which you started using an artificial arm (even in adulthood) would potentially have an effect on reaching accuracy. So, the younger you were when you learned to use an artificial arm the better. However, we found no such correlation between artificial arm errors in amputees and age at first artificial arm use (*r*_*s(12)*_=-0.23, *p*=0.43).

An alternative parallel to the age at first artificial arm use in amputees is the amount of time an individual has spent being limbless before starting to use an artificial arm. So, the longer one waits after amputation to start using an artificial arm, the bigger their reaching errors would be. Here, we find a significant positive correlation between artificial arm absolute errors and years of limbless experience (*r*_*s(12)*_=0.71, *p*=0.005). The sooner an amputee was fitted with an artificial arm after amputation, the smaller errors they made with their current artificial arm years later. Similarly to one-handers, this relationship appears to be driven by motor noise and not by bias. Motor noise was significantly correlated with years of limbless experience (*r*_*s(12)*_*=*0.85, *p*_*bonf*_=0.03), while bias did not (*r*_*s(12)*_=0.32, *p*_*bonf*_=1; comparing between correlations: *Z*=2.9, *p*=0.003). This suggests that the link between age of first use and errors in one-handers may not be limited to a developmental period, but to an individual’s first experiences as a limbless individual. Alternatively, this finding points towards a possible plasticity window in the time after amputation, where early exposure to an artificial arm results in higher levels of control. Although the type of plasticity bottlenecks in each group might be different, it appears that the amount of time an individual spends using their residual limb before starting to use an artificial arm has a long-lasting effect, in terms of motor noise, on their ability to control an artificial arm.

## Discussion

While infants’ reaches are surprisingly functional (Babinsky et al., 2012; von Hofsten, 1980), they take a considerable amount of time to be fine-tuned. There are multiple, non-linear (Olivier et al., 2007) developmental processes occurring until at least the age of ∼12 years (Schneiberg et al., 2002; Simon-Martinez et al., 2018; Sveistrup et al., 2008). In this context, it may not be surprising that we found one-handers, who started using an artificial arm in early childhood, to perform differently in a visuomotor precision reaching task, relative to acquired amputees, who only began to use an artificial arm in adulthood (age *range*=19-56, *mean*=32). Yet, contrary to our expectation, one-handers performed worse than amputees, who in turn did not show any deficits in controlling their artificial arm relative to two-handed controls using their nondominant arm. Considering the early artificial arm experience of one-handers, and that it coincides with a time in development when the motor system has to constantly adapt as the body (and arms) grow, it is surprising that one-handers under-perform at this straightforward motor task relative to amputees. The observed difference between the two artificial arm user’s groups, and the fact that increased experience with an artificial arm did not lead to better performance, is also in stark contrast with the notion that our internal model flexibly scales with the endpoint of the tool, as we use it (Miller et al., 2018).

What mechanisms might be driving the observed deficits in one-handers? To successfully position the artificial arm at the visual target, multiple internal calculations and transformations need to occur, each of which could potentially be impacted by early life experience. First, an internal model of the artificial arm needs to be developed or adapted, so as to translate a desired goal into an appropriate motor coordination plan. Our analysis points at potential deficits in this internal model, as one-handers’ initial directional error–reflecting the execution of an initial motor plan and thought to precede sensory feedback–is greater than that of amputees. If true, this suggests that one-handers may not be able to refine their model enough to create an accurate template of their artificial arm. However, since this deficit was dissociated from our key measure of absolute reaching error, we believe other mechanistic deficits might be at play. With that in mind, a further step for executing a successful reach is being able to integrate concurrent input from the executed plan with online visual feedback of the artificial arm, as well as any other relevant somatosensory feedback from the residual arm. We did not find strong evidence that both static or dynamic localisation of artificial arm position is impaired in one-handers. As such, by elimination, our evidence suggests that it is the process of integrating visual information that is suboptimal in one-handers. This idea is consistent with previous evidence showing one-handers have impaired processing of visual hand information (Maimon-Mor, Schone, et al., 2020). This interpretation is also compatible with previous studies in individuals experiencing a brief period of postnatal visual deprivation which caused long-lasting (though mild) alterations to visuo-auditory processing (de Heering et al., 2016). While the maturation of the vision system occurs much earlier than that of motor control, the ability to optimally integrate visual information continues to develop way into childhood (Contreras-Vidal, 2006; Contreras-Vidal et al., 2005). Lack of concurrent visual and motor experience during development might therefore cause a deficit in the ability to form the computational substrates and thus to integrate visuomotor information. Indeed, we found that endpoint noise, and not initial directional error, associates with age at first artificial arm use. This, too, supports the idea that one’s ability to efficiently integrate sensory information with motor control might relate to early life experience with an artificial arm.

Perhaps our most intriguing result relates to a relationship between the deficits in motor control (reaching error) and the age in infancy at which one-handers started using an artificial arm. Individuals who started using an artificial arm for the first time earlier in infancy also showed less motor deficit. The detected relationship between early life development and motor skill in adulthood allows us to address questions about the plasticity of visuomotor control across life. Why would a 4-year-old child have a disadvantage in visuomotor learning relative to a 2-year-old? This could be explained both by how early she picked up artificial arm use, or rather, how late she waited before starting to use it. While the two alternatives sound similar, these two complementary explanations can be mechanistically dissociated. The first explanation is that the more experience you have with an artificial arm in early childhood, the better you will be at controlling the artificial arm as an adult. Based on the well-accepted assumption that the brain is more plastic early in life, this will allow children to acquire the new skill more easily (Knudsen, 2004). Alternatively, templates for motor control of the hand (e.g., driven by genetics) will decay over time if not consolidated by relevant experience-related input (Dempsey-Jones, Wesselink, et al., 2019; Krubitzer & Prescott, 2018). The longer one waits before including the artificial arm as part of their motor repertoire, the less she will be able to take advantage of this genetic blueprint, i.e. in terms of brain structure and function (Sur & Rubenstein, 2005). Another, third, explanation relates to the fact that by not wearing an artificial arm, one-handers develop alternative strategies to compensate for their missing hand, for example by using their residual-limb. One-handers are known to be proficient residual-limb users, and our previous research shows that the residual arm benefits from the sensorimotor territory normally devoted to the hand (Hahamy et al., 2017; Makin et al., 2013). We also previously showed that residual-limb use in one-handers impacts larger-scale network organisation in sensorimotor cortex (Hahamy et al., 2015) demonstrating how compensatory strategies can affect neural connectivity and dynamics. In an extreme scenario, using the residual-limb as an effector in early life might anchor it as the reference frame for all upper-limb motor control. We previously found that this has implications on peri-personal space representation in one-handers (Maimon-Mor et al., 2017). Thus, the later one-handers start wearing their artificial arm, the later they start developing an alternative reference frame, that is, learning the computations and transformations required to perform movements with an end effector at the artificial arm tip instead of the residual-limb. Another way to think about this competition between alternative strategies is through the prism of ‘habits’ and the idea that once you have perfected a particular motor solution, it is more difficult to update it to a different strategy. As these multiple mechanisms are not mutually exclusive, it is possible that they all contribute to the observed relationship between age at first artificial arm use and reaching errors. Regardless of the specific mechanism, if one-handers sensorimotor processes are optimised for treating the tip of the residual-limb as the end effector, then when wearing an artificial arm, they constantly required to transform information from the residual-limb tip to the artificial arm tip and vice-versa, for extrapolating where the residual-limb needs to be to get the artificial arm tip to a certain target. Similar to skill acquisition of tools, this extra step of transforming information between two spatially removed endpoints may introduce additional noise (integration over space and time).

Amputees, who were born with a complete upper-limb and lost it later in life, did not show similar deficits in artificial arm reaching control. Following the rationale outlined above, it stands to reason that having developed an internal model of their arm in childhood, amputees are able to recycle their internal model of their missing arm to accurately control the artificial arm. Indeed, there is mounting evidence to suggest that amputees maintain the representation of their missing hand long after amputation (Kikkert et al., 2016; Wesselink et al., 2019). The motor requirements for control over an artificial arm are not identical to that of controlling a biological arm, however, from the perspective of the spatial location of the endpoint, the artificial arm roughly mimics that of the missing arm. Interestingly, despite not showing a group deficit in control of their artificial arm (relative to controls’ nondominant arm), we still found that the time they have taken to use the artificial arm following their amputation co-varies with artificial arm reaching errors. As with the one-handers, this relationship could be explained both by how early one picks up artificial arm use, or how late she waits before starting to use it. For example, research in stroke patients suggested that the imbalance triggered by the assault to the brain tissue creates conditions that are favourable for plasticity, and even been referred to as a second sensitive period (Zeiler & Krakauer, 2013). According to this notion, rehabilitation will be most effective within this limited period of plasticity. Similarly, it has been suggested that sensory deprivation can also promote plasticity and learning (Dempsey-Jones, Themistocleous, et al., 2019). Therefore, amputation might cause a cascade that will result in a brief period of increased plasticity that will be more favourable for learning to use an artificial arm. Alternatively, we can also consider the competition model described above; if one has already formed a motor strategy (or a habit) for how to perform tasks without the artificial arm, this learnt strategy will impact her ability to use the artificial arm later on in life. While the reported correlation relies on a smaller sample size and thus should be taken with a little more caution, the fact that overall, amputees do not show systematic deficit relative to controls, indicate that early life experience drives the observed visuomotor deficits in reaching reported here.

To conclude, the fact that one-handers show inferior artificial arm performance compared to amputees is surprising considering both the vast capacity for motor learning that humans exhibit and the fact that one-handers have had more experience with an artificial arm and from a much younger age. By the process of elimination, we have nominated visuomotor integration to be the most likely cause underlying this motor deficit. However, more work testing this interpretation directly needs to be carried out to consolidate our interpretation. Moreover, we found that early life experience with an artificial arm (in the case of one-handers) or early artificial arm experience following amputation (in the case of amputees) has a measurable effect on artificial arm motor precision in adulthood. In our dataset, early life experience with an artificial arm was not a good indicator for successful current artificial arm adoption, however our limited sample size and inclusion criteria of only including individuals who currently use an artificial arm prevents us for making direct clinical recommendations. The relationship between intact-arm performance and current artificial arm use in both one-handers and amputees is also of interest to artificial arm rehabilitation and should be taken into account in future studies. In addition, our research provides insight about the neurocognitive bottlenecks that need to be considered when developing future assistive and augmentative technologies.

## Methods

### Participants

Forty-four artificial arm users were recruited for this study: 21 unilateral acquired amputees (mean age±std = 48.67±12.9, 18 males, 12 with intact right arm), and 23 individuals with congenital unilateral upper-limb loss (transverse deficiency; age ± std = 46.09±11.22, 11 males, 17 with intact right arm; see Table 1 for full demographic details). Sample size was based on recruitment capacities considering the unique populations we tested. Seven participants were excluded from all analyses: 4 participants with a trans-humeral limb-loss (3 amputees) were not able to perform the tasks with their artificial arm. Two participants (1 amputee) had trouble controlling the robotic handle with their artificial arm therefore their artificial arm reaches data in both tasks has been excluded. Data from all tasks involving the robotic manipulandum of one participant (one-hander) was excluded, due to technical issues with the robotic device.

**Table 1.** Demographic details of all participants. Participant: AA = acquired amputee, AC = one-hander, CO = two-handed control; participants marked with an asterisk have valid data only for their intact-hand and were therefore excluded from most analyses. Y since amp = years since amputation. Gender: M = male, F = female. Amp Side = side of limb loss or non-dominant side: L = left, R = right. Amp level = level of limb loss: TR = trans-radial, TH = trans-humeral. Artificial arm type = preferred type of artificial arm: Cos = cosmetic, Mech = mechanical, Myo = myo-electric. Artificial arm wear time = typical number of hours artificial arm worn per week. PAL = functional ability with an artificial arm as determined by PAL questionnaire (0 = minimum function, 1 = maximum function). Usage Score = Artificial arm usage score combining wear time and PAL. Age at first artificial arm use = Age at which one-handers were first introduced to an artificial arm. Years of limbless experience = Time after amputation at which amputees were first introduced to an artificial arm. Residual-limb length = measured in cm.

| Participant | Age | Y Since Amp | Gender | Amp Side | Amp level | Amp cause | Artificial arm Type | Artificial arm wear time | PAL | Usage Score | Age at first artificial arm use | Years of limbless experience | Residual-limb length |
| --- | --- | --- | --- | --- | --- | --- | --- | --- | --- | --- | --- | --- | --- |
| AA01 | 58 | 14 | M | L | TR | Trauma | Myo | 119 | 0.5 | 1.87 |  | 0.5 | 15 |
| AA02 | 46 | 16 | F | L | TR | Trauma | Myo | 56 | 0.59 | 0.49 |  | 0.5 | 14 |
| AA03* | 50 | 3 | F | L | TR | Trauma | Mech | 77 | 0.44 | 0.44 |  |  | 18 |
| AA04* | 53 | 34 | M | L | TH | Trauma | Mech | 48 | 0.2 | -1.4 |  | 0.33 |  |
| AA05 | 21 | 1 | M | R | TR | Trauma | None | 0 | 0.04 | -3.44 |  | 0.083 | 34 |
| AA06* | 42 | 18 | M | R | TR | Trauma | Cos | 35 | 0.07 | -2.33 |  |  | 16 |
| AA07 | 61 | 21 | M | L | TR | Trauma | Cos | 105 | 0.67 | 2.21 |  | 0.125 | 29.5 |
| AA08 | 60 | 42 | M | R | TR | Trauma | Mech | 87.5 | 0.28 | 0.05 |  | 3.5 | 9 |
| AA09* | 65 | 37 | M | R | TH | Trauma | Mech | 98 | 0.46 | 1.11 |  | 0.25 |  |
| AA10* | 47 | 21 | M | R | TH | Trauma | Cos | 84 | 0.3 | 0.03 |  | 1 |  |
| AA11 | 68 | 12 | M | L | TR | Trauma | Mech | 35 | 0.54 | -0.31 |  | 0.33 | 21.5 |
| AA12 | 49 | 5 | M | R | TR | Vascular disease | Cos | 42 | 0.59 | 0.1 |  | 0.5 | 20 |
| AA13 | 57 | 29 | M | L | TR | Trauma | Mech | 65 | 0.11 | -1.31 |  | 1 | 18 |
| AA14 | 53 | 33 | M | L | TR | Trauma | Myo | 98 | 0.43 | 0.98 |  | 0.67 | 12 |
| AA15* | 28 | 10 | F | R | TR | Trauma | Mech | 2 | 0 | -3.55 |  | 5 | 7.5 |
| AA16 | 29 | 11 | M | L | TR | Trauma | None | 0 | 0 | -3.61 |  | 2 | 28.5 |
| AA17 | 43 | 20 | M | R | TR | Trauma | Myo | 98 | 0.61 | 1.75 |  | 0.083 | 8 |
| AA18 | 55 | 12 | M | L | TR | Trauma | Myo | 98 | 0.65 | 1.92 |  | 0.5 | 33 |
| AA19 | 61 | 17 | M | L | TR | Trauma | Mech | 91 | 0.74 | 2.11 |  | 1 | 18 |
| AA21 | 30 | 3 | M | L | TR | Trauma | Myo | 49 | 0.59 | 0.29 |  | 0.54 | 20.5 |
| AC01 | 51 |  | F | L | TR | Congenital | Cos | 7 | 0.26 | -2.3 | 0.5 |  | 10 |
| AC02 | 47 |  | M | L | TR | Congenital | Mech | 84 | 0.7 | 1.75 | 1 |  | 13 |
| AC03 | 45 |  | F | L | TR | Congenital | Myo | 63 | 0.46 | 0.13 | 4.5 |  | 8 |
| AC04* | 26 |  | M | L | TR | Congenital | Mech | 6 | 0.13 | -2.88 | 0.25 |  | 15 |
| AC05* | 55 |  | F | L | TR | Congenital | Cos | 112 | 0.3 | 0.82 | 0.25 |  | 6 |
| AC06 | 63 |  | M | L | TR | Congenital | Cos | 87.5 | 0.35 | 0.35 | 2 |  | 10 |
| AC07 | 35 |  | M | L | TR | Congenital | Cos | 56 | 0.28 | -0.84 | 3 |  | 11 |
| AC09 | 49 |  | M | L | TR | Congenital | Myo | 91 | 0.57 | 1.39 | 2 |  | 21 |
| AC10 | 42 |  | M | L | TR | Congenital | Cos | 56 | 0.54 | 0.28 | 2 |  | 10.5 |
| AC11 | 66 |  | F | R | TR | Congenital | Cos | 42 | 0.35 | -0.93 | 5 |  | 9 |
| AC12 | 56 |  | F | R | TR | Congenital | Cos | 98 | 0.43 | 0.98 | 3 |  | 11.5 |
| AC13* | 53 |  | M | L | TH | Congenital | Mech | 63 | 0.33 | -0.43 | 2 |  |  |
| AC14 | 42 |  | M | L | TR | Congenital | Mech | 2 | 0.09 | -3.17 | 4 |  | 12 |
| AC15 | 55 |  | F | L | TR | Congenital | Myo | 105 | 0.65 | 2.12 | 3 |  | 11.5 |
| AC17 | 29 |  | M | L | TR | Congenital | Myo | 70 | 0.46 | 0.33 | 1 |  | 12 |
| AC18 | 53 |  | F | L | TR | Congenital | Cos | 48 | 0.65 | 0.52 | 1.5 |  | 7 |
| AC20 | 52 |  | F | R | TR | Congenital | Myo | 32.5 | 0.26 | -1.58 | 0.3 |  | 11.5 |
| AC21 | 32 |  | F | R | TR | Congenital | Myo | 40 | 0.41 | -0.73 | 0.5 |  | 9 |
| AC22 | 57 |  | M | R | TR | Congenital | Mech | 126 | 0.69 | 2.88 | 2 |  | 15.5 |
| AC23 | 47 |  | F | L | TR | Congenital | Myo | 84 | 0.89 | 2.56 | 11 |  | 8.5 |
| AC25 | 41 |  | M | L | TR | Congenital | Myo | 112 | 0.85 | 3.17 | 0.25 |  | 8 |
| CO01 | 28 |  | F | L |  |  |  |  |  |  |  |  |  |
| CO03 | 40 |  | F | L |  |  |  |  |  |  |  |  |  |
| CO04 | 59 |  | M | L |  |  |  |  |  |  |  |  |  |
| CO05 | 27 |  | F | R |  |  |  |  |  |  |  |  |  |
| CO06 | 61 |  | F | L |  |  |  |  |  |  |  |  |  |
| CO07 | 35 |  | M | L |  |  |  |  |  |  |  |  |  |
| CO08 | 34 |  | F | L |  |  |  |  |  |  |  |  |  |
| CO09 | 24 |  | M | L |  |  |  |  |  |  |  |  |  |
| CO10* | 70 |  | M | L |  |  |  |  |  |  |  |  |  |
| CO11 | 24 |  | M | L |  |  |  |  |  |  |  |  |  |
| CO12 | 18 |  | F | L |  |  |  |  |  |  |  |  |  |
| CO13 | 67 |  | M | L |  |  |  |  |  |  |  |  |  |
| CO14 | 50 |  | M | R |  |  |  |  |  |  |  |  |  |
| CO15 | 51 |  | F | L |  |  |  |  |  |  |  |  |  |
| CO16 | 36 |  | F | L |  |  |  |  |  |  |  |  |  |
| CO17 | 41 |  | M | R |  |  |  |  |  |  |  |  |  |
| CO18 | 33 |  | M | R |  |  |  |  |  |  |  |  |  |
| CO19 | 45 |  | M | R |  |  |  |  |  |  |  |  |  |
| CO20 | 54 |  | M | R |  |  |  |  |  |  |  |  |  |
| CO21 | 53 |  | M | L |  |  |  |  |  |  |  |  |  |

Additionally, twenty age, gender, and handedness matched two-handed controls were recruited for this study (mean age ± std = 42.55 ± 15.5, 11 males, 14 with a dominant right arm). For all analyses, the controls’ dominant arm was compared to the intact arm of the artificial arm users, and the non-dominant arm was compared to the artificial arm. For the sake of brevity, we refer to the dominant arm of controls as the intact-hand and the non-dominant arm as the artificial arm.

Participants were recruited to the study between October 2017 and December 2018, based on the guidelines in our ethical approvals (UCL REC: 9937/001; NHS National Research Ethics service: 18/LO/0474), and in accordance with the declaration of Helsinki. The following inclusion criteria were taken into consideration during recruitment: (1) 18 to 70 years old, (2) MRI safe (for the purpose of other tasks conducted in the scanner), (3) no previous history of mental disorders, (4) for one-handers, owned at least one type of prosthesis during recruitment, (5) for acquired amputees, amputation occurred at least 6 months before recruitment. All participants gave full written informed consent for their participation, data storage and dissemination.

### Main Task

#### Experimental setup

Participants sat in front of the experimental apparatus on a barber-style chair, with their head leaning against a forehead rest. Participants performed horizontal plane reaches while holding a handle of a robotic manipulandum with either their hand or the artificial arm, with the arm strapped to an armrest. A monitor displaying the task was viewed via a mirror, such that participants did not have direct vision of their arm. To further block any vision of the participant’s limb a black barber’s cape was used to cover their entire upper body, including their elbow and shoulder. Continuous visual feedback of the robotic handle’s position (i.e., the intact/artificial arm position) was delivered as a 4 *cm* diameter white cursor (representing the handle size) with a 0.3 *cm* diameter circle at the centre. The handle’s position was recorded with a sampling frequency of 200 *Hz*.

### Experimental design – main task

Participants were asked to reach to visual targets while receiving visual feedback of their hand position using each of their arms. To ensure the setup was optimised for artificial arm reaches, participants performed the task with their non-dominant/artificial arm first. At the beginning of each trial set 6 practice trials (using targets not included in the task target set) were presented to the participant. Practice was repeated until both experimenter and participant were happy that the participant felt comfortable and the instructions were fully understood.

A trial was initiated once participants placed the cursor within a white square (1.5 *cm* × 1.5 *cm*) indicating the home (start) position (denoted as position [0,0]). Participants were situated so the home position was aligned with their midline. In each trial, participants reached to a visual target (1.5 *cm* × 1.5 *cm* square) presented in one of 60 predefined locations (see Figure 1B). At the end of a trial, the target changed colour to blue to indicate the reach has ended and the endpoint position was recorded. To reduce fatigue and experiment duration, participants were then mechanically assisted by the manipulandum moving the handle back to the home position, before the start of the next trial.

To quantify participants’ biological and artificial arm motor control, participants were asked to perform a single movement to the target and avoid corrective movements. Constant visual feedback of the arm’s position was given. To encourage participants to perform fast-reaching movements, a maximum movement time of 1 *sec* per reach was imposed. Movement initiation was defined by arm velocity exceeding 3.5 *cm/s* starting from the time of participants first movement, following the presentation of the target.

#### Data processing and analysis – main task

To identify the end of the first reach in each trial, tangential arm velocities were used to determine movement termination. Velocity data were smoothed using an 8 Hz low-pass Butterworth filter (Przybyla et al., 2013). Movement termination was defined by the first minimum with a velocity smaller than 50% of the peak velocity. We note that very similar results were observed when using the end of the trial (1 *sec* after initiation) as the movement termination time. Individual trials were excluded if they were accidentally initiated, i.e., if movement terminated close to the home position – closer than 2 *cm* or with a y-value (depth) smaller than 1 *cm*; or if the participants did not finish their reach at the end of the allocated time (1 *sec*) – i.e., trials where movement >10 *cm/s* was recorded at the end of the trial. An average of 1.1 and 1.4 trials per participant were excluded for the intact and artificial arm respectively, with a range of 0-7. There were no group differences in the amount of excluded trials. Artificial arm reach data from one amputee and one one-hander were excluded, due to technical issues with the device.

Absolute Euclidean error from the target was used as the main measure (See Figure 1A&C). Motor noise (variability) and bias were calculated for each participant, for each arm, by aggregating across all targets. Error vectors of each trial (the line connecting the target with the end-point location) were overlaid as if they were made to a single target (See Figure 2A). Bias was defined as the distance from the centre of the overlaid end-points (calculated as the mean x and mean y of the relative error vectors) to the target. Noise was defined as the spatial standard deviation (SD) of endpoints relative to the same centre of overlaid endpoints. Initial directional error was defined for each trial as the absolute angle between the direction vector–the line connecting the home position and the arm’s location at peak velocity (see Figure 2B); and the direction vector–the line connecting the target and the home position. The arm’s location at peak velocity was used as a proxy for a time-point that mostly reflects the motor planning phase, i.e. feedforward mechanisms, before sensory information is used to correct for errors during execution.

### Additional Tasks

#### 1D arm localisation

To assess residual-limb and artificial arm sense of limb-position, we used a task similar to that described in (McDonnell et al., 1989). Participants were asked to place their residual-limb or artificial arm in an opaque tube (see Figure 2C). In each trial, an adjustable contact plate was placed at a different position within the tube. Participants were asked to move their arm into the tube until it made contact with the plate. They were then instructed to use their intact arm to mark the estimated location of their tested arm on a paper strip placed on the side of the tube. At the end of the trial, participants were asked to remove their arm from the tube before the start of the next trial. A barber’s cape was used to cover the upper body and arms. For each condition (residual/artificial arm), pseudo-randomised 8 distances were used; these were calculated as a percentage of participant’s maximum reach distance (25%-75%). The mean absolute distance between the participant’s estimate and true position was used as a measure of 1D localisation abilities. Two amputees and two one-handers did not take part in this task as it was introduced later in the data collection process.

#### 2D arm localisation (reaching without visual feedback)

The 2D arm localisation task was almost identical to our main task, with the exception that participants reached to visual targets without receiving continuous visual feedback of their limb position and were allowed to perform corrective movements. To prevent a preceptive drift from the lack of visual information of limb position, visual feedback of the arm’s position was given at the end of the trial when returning to the start position. The cursor only reappeared when the arm was less than 3 *cm* away from the start position. At the beginning of the trial, the cursor disappeared upon movement initiation. Movement termination and the recording of the arm’s final position occurred when velocity was less than 3.5 *cm/s* for more than 1 *sec*. Due to the noisier nature of these reaches, each of the 60 targets used in the main task was repeated twice (i.e., a total of 120 trials for each arm).

Absolute Euclidean error from the target was used as the main measure. Individual trials were excluded if they were accidentally initiated (see main task data analysis protocol); or if the trial was suspected as invalid, i.e., movement time was longer than 10 secs or error was larger than 20 cm. An average of 1.35 and 1.5 trials per participant were excluded for the intact and artificial arm respectively, with a range of 0-13. There were no group differences in the amount of excluded trials. Two participants only produced partial data, missing artificial arm reaches of 1 one-hander and dominant arm reaches of 1 control.

### Artificial arm usage assessment

Participants completed a questionnaire to assess various aspects of their current and past artificial arm use. Frequency and functionality of artificial arm use were combined to create an overall artificial arm usage score (as previously used in (Maimon-Mor, Obasi, et al., 2020; Maimon-Mor & Makin, 2020; van den Heiligenberg et al., 2017; Van Den Heiligenberg et al., 2018). To determine frequency of use, participants were asked to indicate the typical number of hours per day, and days per week, they use their artificial arm. These were then used to determine the typical number of hours per week that the artificial arm was worn. To determine functionality of artificial arm use, participants were asked to complete the artificial arm activity log (Prosthesis Activity Log - PAL), a modified version of the Motor Activity Log (MAL) questionnaire, which is commonly used to assess arm functionality in those with upper-limb impairments (Uswatte et al., 2006). The PAL consists of a list of 27 daily activities (see https://osf.io/jfme8/); participants rated how often they incorporate their artificial arm to complete each activity on a scale of “never” (0 points), “sometimes” (1 point) or “very often” (2 points). The PAL score is then calculated as the sum of all points divided by the maximum possible score, generating a value between 0 (no functionality) and 1 (maximum functionality). Artificial arm scores were calculated for the most used artificial arm, wear time and PAL were standardised using a Z-transform and summed to create an artificial arm usage score that reflects frequency of use and incorporation of the artificial arm in activities of daily living. These measures have been previously shown to have good reliability using a test-retest assessment (Maimon-Mor, Obasi, et al., 2020).

One-handers were asked to report the age at which they first used an artificial arm. Two participants (AC16, AC17) were assigned a value of one year after responding: “months old” and “less than a year” respectively. Acquired amputees were asked to report the time after amputation in which they were fitted with their first artificial arm. Two participants (AA13, AA19) were assigned a value of one year after responding: “same year, few months after amputation” and “A few months after amputation” respectively.

### Statistical analyses

Statistical analyses were carried out using JASP (Jasp Team, 2020) An analysis of covariance (ANCOVA) was used to test group differences for all measures in which we had recorded performance of both arms (i.e., all measures but 1D localisation). The artificial arm performance was the dependent variable, intact-hand performance was defined as a covariate and group (controls, amputees, and one-handers) as a between-subject variable. Post-hoc group tests were corrected for multiple comparisons (Tukey correction). Absolute error measures were logarithmically transformed and then averaged in order to correct for the skewed error distribution and satisfy the conditions for parametric statistical testing. Outliers were defined as 1.5 times the IQR (interquartile ranges) below the first quartile or above the third quartile of the transformed error. Following this outlier criteria, in the main task, 2 participants (1 amputee, 1 one-hander) were excluded due to their high artificial arm errors. For the 2D localisation task, 3 participants (2 amputees, 1 control) were excluded due to their high intact-arm errors. In parametric analyses (ANCOVA, ANOVA, Pearson correlations), where the frequentist approach yielded a non-significant p-value, a parallel Bayesian approach was used and Bayes Factors (BF) were reported (Morey & Rouder, 2015; Rouder et al., 2009, 2012, 2016). A BF<0.33 is interpreted as support for the null-hypothesis, BF > 3 is interpreted as support for the alternative hypothesis (Dienes, 2014). In Bayesian ANOVAs and ANCOVA’s the inclusion Bayes Factor of an effect (BF_Incl_) is reported, reflecting that the data is X (BF) times more likely under the models that include the effect than under the models without this predictor. When using a Bayesian t-test, a Cauchy prior width of 1.39 was used, this was based on the effect size of the main task, when comparing artificial arm reaches of amputees and one-handers. Therefore, the null hypothesis in these cases would be there is no effect as large as the effect observed in the main task. Parametric analyses were used if assumptions (e.g. for normality) were met, otherwise a Spearman correlation/Mann-Whitney were used. Since the Spearman correlation has, to our knowledge, no current Bayesian implementation no BF values are reported for these tests. The Parametric Watson-Williams multi-sample test for equal means was used as a one-way ANOVA test for bias angular data.

## Acknowledgments

This work was supported by an ERC Starting Grant (715022 EmbodiedTech), awarded to TRM, who was further funded by a Wellcome Trust Senior Research Fellowship (215575/Z/19/Z). R.O.M.M. is supported by the Clarendon scholarship and University College, Oxford. We thank Nour Odeh, Mischa Dhar and Victoria Root for data collection and Dorothy Cowie and Jordan Taylor for their comments on the manuscript. We thank Opcare for their help in participants recruitment, and our participants and their families for their ongoing support of our research.

## Supplementary Information

### Supplementary Results

#### Speed-accuracy trade-off in artificial arm reaches

To test whether the three groups (controls, amputees and one-handers) use the same speed accuracy trade-off strategy. More specifically, whether artificial arm reaches follow the same control principles as biological arm reaches that have been shown to follow Fitts’ law (Fitts 1954), a specific relationship between the movement time and the movement distance:

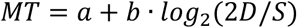

[MT: movement time, D: distance to target, S: target size – constant]

For each subject, we used a linear regression to obtain the parameters *a* and *b*. To test whether reaches of each group follow Fitts’ Law equally, the r-squared value of the regression was compared across all groups as well as the regression parameters (using an ANCOVA, controlling for the parallel measure of the intact hand). To reduce the influence of noisy individual reaches, the reaches have been divided and averaged into 6 bins, based on their distance from the starting position. We found no group differences in either goodness of fit (***r***^***2***^: *p*=0.84, *BF*_*Incl*_=0.167) or fitted parameters (***a***: *p*=0.31, *BF*_*Incl*_=0.347, ***b***: *p=0*.*61, BF*_*Incl*_*=0*.*22*) between groups, indicating artificial arms reaches follow Fitts’ laws and do not differ in their speed-accuracy trade-off strategy (see Figure S1, Table S4 for full statistical reports and https://osf.io/quyke/ for plots of individual participants).

### Movement maximum velocity

For each participant, the maximum velocity of every trial was extracted and averaged across all trials. When comparing the mean velocity between groups (while controlling for the velocity of the intact hand), we found a significant relationship with intact hand velocity (*F*_*(1,47*_)= 237.615, *p*<0.001, η_p_^2^=0.835) and a significant group effect (*F*_*(2,47)*_=3.49, *p*=0.04, η_p_^2^=0.13). Post-hoc comparisons showed artificial arm reaches of amputees were slightly, but not significantly, faster than one-handers (*t*=-2.31, *p*_*tukey*_=0.06, *Cohen’s-d*=-0.31). Amputees were also slightly, but not significantly, faster than controls’ non-dominant hand reaches (*t*=-2.37, *p*_*tukey*_=0.06, *Cohen’s-d*=-0.36). One-handers artificial arm velocities did not differ from those of controls (*t*=0.19, *p*_*tukey*_=0.98, *Cohen’s-d*=0.03). Importantly, these differences in velocities were not related to our main effect of group differences in reaching accuracy. When adding the maximum reaching velocities as a covariate to the main analysis described in section 3.1, all reported results remained significant and the effect of maximum velocity on absolute reaching error was not significant (F(1,46)= 0.27, p=0.61, *BF*_*Incl*_=0.247).

## Supplementary Figures

**Figure S1.**
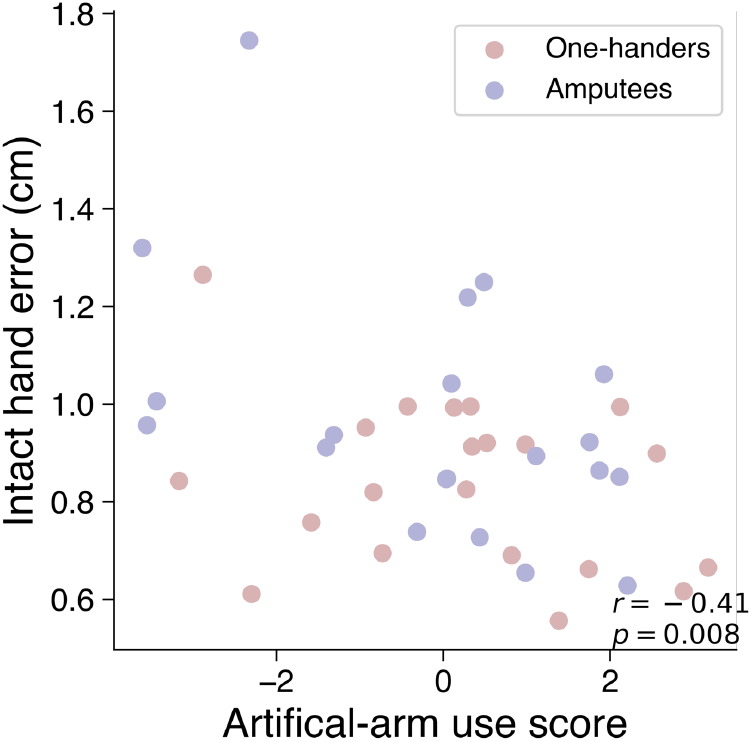
Intact hand errors and daily artificial arm use. We found a significant correlation (*r*_(39)_=-0.41, *p*=0.008) between artificial arm daily use and intact-hand reaching errors. In this analysis, both artificial arm users’ groups (one-handers and amputees) were analysed together as we found no differences in intact-hand errors between the groups. Daily artificial arm use was quantified using questionnaires relating to both wear-time and functionality of use.

**Figure S2.**
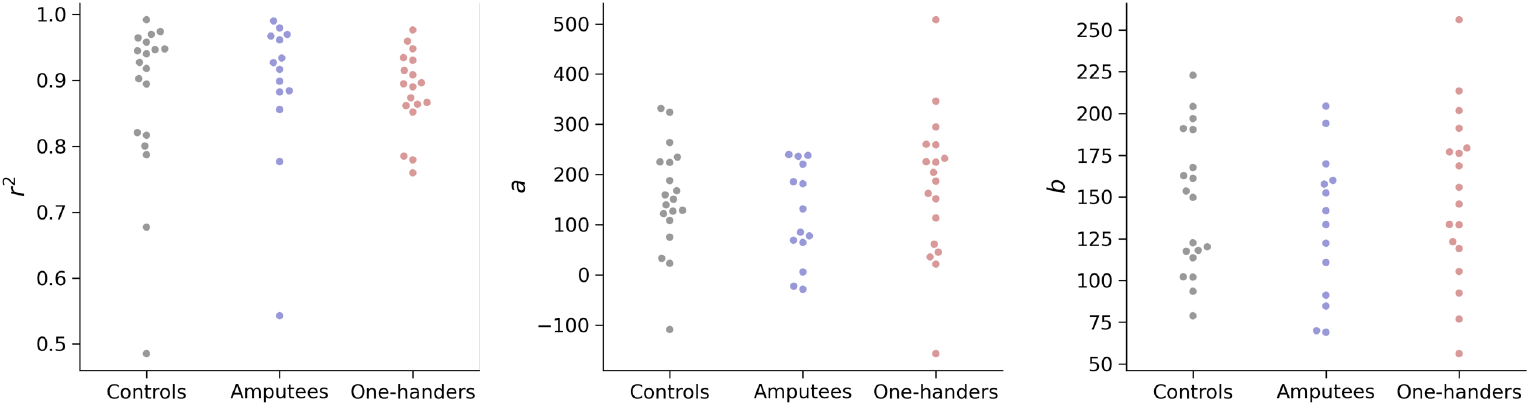
Group values for Fitts law model fitting (r^2^,a,b). A linear regression was fit for each participant’s reaches to obtain the Fitts law model’s parameters *a* and *b*. Parameters, as well as goodness-of-fit (*r*_^*2*^_), were compared across groups. We found no group differences in either goodness of fit (***r***^***2***^: *p*=0.84, *BF*_*Incl*_=0.167) or fitted parameters (***a***: *p*=0.31, *BF*_*Incl*_=0.347, ***b***: *p=0*.*61, BF*_*Incl*_*=0*.*22*) between groups, indicating artificial arms reaches follow Fitts’ laws and do not differ in their speed-accuracy trade-off strategy

## Supplementary Tables

**Table S1.** Main analysis while controlling for artificial arm/nondominant-arm side. Results of a follow-up ANCOVA analysis showing no effects of artificial arm side (L vs R) on artificial arm reaching errors. Our main finding of a significant group effect was also unaffected by accounting for the side of the arm making the reaches.

| Factors | SS | df | MS | F | p |
| --- | --- | --- | --- | --- | --- |
| Group – Fixed factor | 0.719 | 2 | 0.360 | 12.117 | < .001 |
| Artificial arm side – Fixed factor | 0.033 | 1 | 0.033 | 1.119 | 0.296 |
| Intact-arm absolute errors - Covariate | 0.835 | 1 | 0.835 | 28.127 | < .001 |
| Group * Artificial arm side interaction | 0.013 | 2 | 0.006 | 0.216 | 0.807 |
| Residuals | 1.306 | 44 | 0.030 |  |  |

**Table S2.** Main analysis while controlling for residual-limb length. Results of a follow-up ANCOVA analysis showing no effects of residual-limb length on artificial arm reaching errors. Our main finding of a significant group effect was also unaffected by accounting for residual-limb length. Note that this analysis only includes artificial arm users (amputees and one-handers) as controls have a complete arm and therefore no residual-limb length.

| Factors | SS | df | MS | F | p |
| --- | --- | --- | --- | --- | --- |
| Group – Fixed factor | 0.137 | 1 | 0.137 | 5.065 | 0.032 |
| Residual-limb length- Covariate | 0.042 | 1 | 0.042 | 1.565 | 0.221 |
| Intact-arm absolute errors - Covariate | 0.318 | 1 | 0.318 | 11.768 | 0.002 |
| Residuals | 0.758 | 28 | 0.027 |  |  |

**Table S3.** Comparing artificial arm error noise while controlling for artificial arm bias. Results of a follow-up ANCOVA analysis showing that while there is a significant relationship between bias and noise, the group differences in error noise are independent of bias.

| Factors | SS | df | MS | F | p |
| --- | --- | --- | --- | --- | --- |
| Group – Fixed factor | 1.434 | 2 | 0.717 | 12.405 | < .001 |
| Artificial arm bias - Covariate | 0.404 | 1 | 0.404 | 6.991 | 0.011 |
| Intact-arm noise - Covariate | 0.460 | 1 | 0.460 | 7.962 | 0.007 |
| Residuals | 2.659 | 46 | 0.058 |  |  |

**Table S4.**
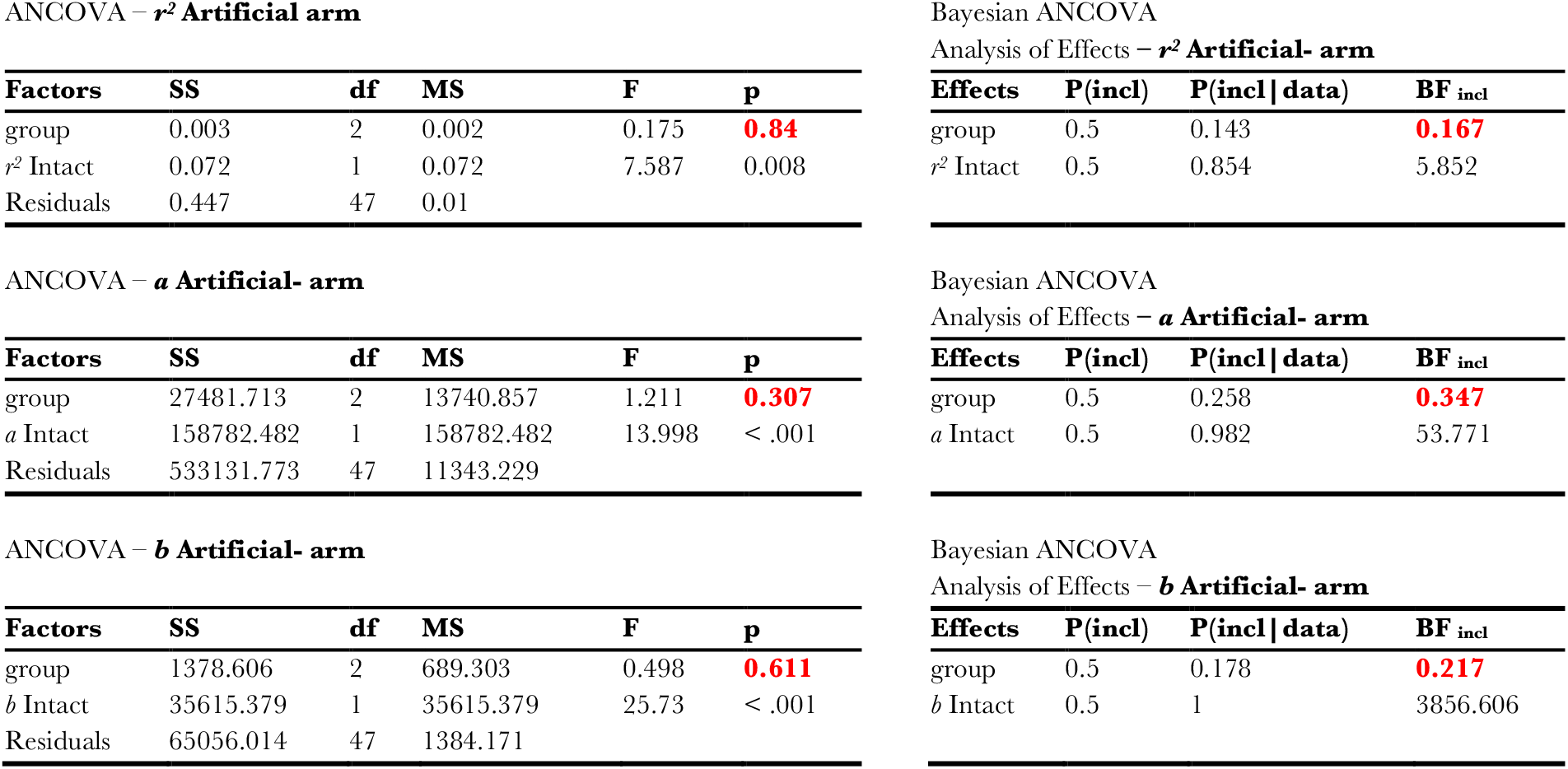
Frequentist and Bayesian analysis of model fitting reaches data to Fits’ Law. Full statistical report of group comparisons of model’s parameters *a* and *b* as well as goodness-of-fit (*r*^*2*^) of the linear regression model. No differences were found across groups.

ANCOVA – $r^2$ Artificial arm
| Factors | SS | df | MS | F | p |
| --- | --- | --- | --- | --- | --- |
| group | 0.003 | 2 | 0.002 | 0.175 | 0.84 |
| $r^2$ Intact | 0.072 | 1 | 0.072 | 7.587 | 0.008 |
| Residuals | 0.447 | 47 | 0.01 |  |  |

ANCOVA – $a$ Artificial- arm
| Factors | SS | df | MS | F | p |
| --- | --- | --- | --- | --- | --- |
| group | 27481.713 | 2 | 13740.857 | 1.211 | 0.307 |
| $a$ Intact | 158782.482 | 1 | 158782.482 | 13.998 | < .001 |
| Residuals | 533131.773 | 47 | 11343.229 |  |  |

ANCOVA – $b$ Artificial- arm
| Factors | SS | df | MS | F | p |
| --- | --- | --- | --- | --- | --- |
| group | 1378.606 | 2 | 689.303 | 0.498 | 0.611 |
| $b$ Intact | 35615.379 | 1 | 35615.379 | 25.73 | < .001 |
| Residuals | 65056.014 | 47 | 1384.171 |  |  |

Bayesian ANCOVA Analysis of Effects – $r^2$ Artificial- arm
| Effects | P(incl) | P(incl \| data) | BF<sub>incl</sub> |
| --- | --- | --- | --- |
| group | 0.5 | 0.143 | 0.167 |
| $r^2$ Intact | 0.5 | 0.854 | 5.852 |

Bayesian ANCOVA Analysis of Effects – $a$ Artificial- arm
| Effects | P(incl) | P(incl \| data) | BF<sub>incl</sub> |
| --- | --- | --- | --- |
| group | 0.5 | 0.258 | 0.347 |
| $a$ Intact | 0.5 | 0.982 | 53.771 |

| Effects | P(incl) | P(incl \| data) | BF<sub>incl</sub> |
| --- | --- | --- | --- |
| group | 0.5 | 0.178 | 0.217 |
| $b$ Intact | 0.5 | 1 | 3856.606 |

## References

Adolph, K. E., & Franchak, J. M. (2017). The development of motor behavior. In Wiley Interdisciplinary Reviews: Cognitive Science (Vol. 8, Issues 1–2). Wiley-Blackwell. https://doi.org/10.1002/wcs.1430

Babinsky, E., Braddick, O., & Atkinson, J. (2012). Infants and adults reaching in the dark. Experimental Brain Research, 217(2), 237–249. https://doi.org/10.1007/s00221-011-2984-5

Berthier, N. E., & Keen, R. (2006). Development of reaching in infancy. Experimental Brain Research, 169(4), 507–518. https://doi.org/10.1007/s00221-005-0169-9

Biddiss, E., & Chau, T. (2007). The roles of predisposing characteristics, established need, and enabling resources on upper extremity prosthesis use and abandonment. Disability & Rehabilitation: Assistive Technology, 2(2), 71–84. https://doi.org/10.1080/17483100601138959

Bruurmijn, M. L. C. M., Pereboom, I. P. L., Vansteensel, M. J., Raemaekers, M. A. H., & Ramsey, N. F. (2017). Preservation of hand movement representation in the sensorimotor areas of amputees. Brain, 140(12), 3166–3178. https://doi.org/10.1093/brain/awx274

Contreras-Vidal, J. L. (2006). Development of forward models for hand localization and movement control in 6-to 10-year-old children. Human Movement Science, 25(4–5), 634–645. https://doi.org/10.1016/j.humov.2006.07.006

Contreras-Vidal, J. L., Bo, J., Boudreau, J. P., & Clark, J. E. (2005). Development of visuomotor representations for hand movement in young children. Experimental Brain Research, 162(2), 155–164. https://doi.org/10.1007/s00221-004-2123-7

de Heering, A., Dormal, G., Pelland, M., Lewis, T., Maurer, D., & Collignon, O. (2016). A Brief Period of Postnatal Visual Deprivation Alters the Balance between Auditory and Visual Attention. Current Biology, 26(22), 3101–3105. https://doi.org/10.1016/j.cub.2016.10.014

Dempsey-Jones, H., Themistocleous, A. C., Carone, D., Ng, T. W. C., Harrar, V., & Makin, T. R. (2019). Blocking Tactile Input to One Finger Using Anaesthetic Enhances Touch Perception and Learning in Other Fingers. Journal of Experimental Psychology: General, 148(4), 713–727. https://doi.org/10.1037/xge0000514.supp

Dempsey-Jones, H., Wesselink, D. B., Friedman, J., & Makin, T. R. (2019). Organized Toe Maps in Extreme Foot Users. Cell Reports, 28(11), 2748-2756.e4. https://doi.org/10.1016/j.celrep.2019.08.027

Dienes, Z. (2014). Using Bayes to get the most out of non-significant results. Frontiers in Psychology, 5(July), 1–17. https://doi.org/10.3389/fpsyg.2014.00781

Elliott, D., Chua, R., & Helsen, W. F. (2001). A century later: Woodworth’s (1899) two-component model of goal-directed aiming. Psychological Bulletin, 127(3), 342–357. https://doi.org/10.1037/0033-2909.127.3.342

Faisal, A. A., Selen, L. P. J., & Wolpert, D. M. (2008). Noise in the nervous system. Nature Reviews Neuroscience, 9(april), 292–303. https://doi.org/10.1038/nrn2258

Gandhi, T. K., Singh, A. K., Swami, P., Ganesh, S., & Sinha, P. (2017). Emergence of categorical face perception after extended early-onset blindness. Proceedings of the National Academy of Sciences of the United States of America, 114(23), 6139–6143. https://doi.org/10.1073/pnas.1616050114

Gordon, J., Ghilardi, M. F., & Ghez, C. (1995). Impairments of reaching movements in patients without proprioception. I. Spatial errors. Journal of Neurophysiology, 73(1), 347–360. https://doi.org/10.1152/jn.1995.73.1.347

Hahamy, A., Macdonald, S. N., van den Heiligenberg, F. M. Z., Kieliba, P., Emir, U., Malach, R., Johansen-Berg, H., Brugger, P., Culham, J. C., & Makin, T. R. (2017). Representation of Multiple Body Parts in the Missing-Hand Territory of Congenital One-Handers. Current Biology, 27(9), 1350–1355. https://doi.org/10.1016/j.cub.2017.03.053

Hahamy, A., Sotiropoulos, S. N., Henderson Slater, D., Malach, R., Johansen-Berg, H., & Makin, T. R. (2015). Normalisation of brain connectivity through compensatory behaviour, despite congenital hand absence. ELife, 4. https://doi.org/10.7554/eLife.04605.

Jasp Team. (2020). JASP (0.12.2).

Karmiloff-Smith, A. (1998). Development itself is the key to understanding developmental disorders. In Trends in Cognitive Sciences (Vol. 2, Issue 10, pp. 389–398). Elsevier Current Trends. https://doi.org/10.1016/S1364-6613(98)01230-3

Kikkert, S., Kolasinski, J., Jbabdi, S., Tracey, I., Beckmann, C. F., Johansen-Berg, H., & Makin, T. R. (2016). Revealing the neural fingerprints of a missing hand. ELife, 5. https://doi.org/10.7554/eLife.15292

Knudsen, E. I. (2004). Sensitive periods in the development of the brain and behavior. Journal of Cognitive Neuroscience, 16(8), 1412–1425. https://doi.org/10.1162/0898929042304796

Krakauer, J. W., Ghilardi, M. F., & Ghez, C. (1999). Independent learning of internal models for kinematic and dynamic control of reaching. Nature Neuroscience, 2(11), 1026–1031. https://doi.org/10.1038/14826

Krubitzer, L. A., & Prescott, T. J. (2018). Special Issue: Time in the Brain The Combinatorial Creature: Cortical Phenotypes within and across Lifetimes. Trends in Neurosciences, 41, 744–762. https://doi.org/10.1016/j.tins.2018.08.002

Leed, J. E., Chinn, L. K., & Lockman, J. J. (2019). Reaching to the Self: The Development of Infants’ Ability to Localize Targets on the Body. Psychological Science, 30(7), 1063–1073. https://doi.org/10.1177/0956797619850168

Maimon-Mor, R. O., Johansen-Berg, H., & Makin, T. R. (2017). Peri-hand space representation in the absence of a hand – Evidence from congenital one-handers. Cortex, 95, 169–171. https://doi.org/10.1016/j.cortex.2017.08.016

Maimon-Mor, R. O., & Makin, T. R. (2020). Is an artificial limb embodied as a hand? Brain decoding in prosthetic limb users. PLOS Biology, 18(6), e3000729. https://doi.org/10.1371/journal.pbio.3000729

Maimon-Mor, R. O., Obasi, E., Lu, J., Odeh, N., Kirker, S., MacSweeney, M., Goldin-Meadow, S., & Makin, T. R. (2020). Talking with Your (Artificial) Hands: Communicative Hand Gestures as an Implicit Measure of Embodiment. IScience, 23(11). https://doi.org/10.1016/j.isci.2020.101650

Maimon-Mor, R. O., Schone, H. R., Moran, R., Brugger, P., & Makin, T. R. (2020). Motor control drives visual bodily judgements. Cognition, 196. https://doi.org/10.1016/j.cognition.2019.104120

Makin, T. R., Cramer, A. O., Scholz, J., Hahamy, A., Henderson Slater, D., Tracey, I., & Johansen-Berg, H. (2013). Deprivation-related and use-dependent plasticity go hand in hand. ELife, 2013, 1–15. https://doi.org/10.7554/eLife.01273.01273

McDonnell, P. M., Scott, R. N., Dickison, J., Anne Theriault, R., & Wood, B. (1989). Do artificial limbs become part of the user? New evidence. Journal of Rehabilitation Research and Development, 26(2), 17– 24.

Miller, L. E., Montroni, L., Koun, E., Salemme, R., Hayward, V., & Farnè, A. (2018). Sensing with tools extends somatosensory processing beyond the body. Nature, 561(7722), 239–242. https://doi.org/10.1038/s41586-018-0460-0

Miyashita-Lin, E. M., Hevner, R., Wassarman, K. M., Martinez, S., & Rubenstein, J. L. R. (1999). Early neocortical regionalization in the absence of thalamic innervation. Science, 285(5429), 906– 909. https://doi.org/10.1126/science.285.5429.906

Morey, R. D., & Rouder, J. N. (2015). BayesFactor (0.9.10-2).

Olivier, I., Hay, L., Bard, C., & Fleury, M. (2007). Age-related differences in the reaching and grasping coordination in children: Unimanual and bimanual tasks. Experimental Brain Research, 179(1), 17– 27. https://doi.org/10.1007/s00221-006-0762-6

Patterson, J. R., Brown, L. E., Wagstaff, D. A., & Sainburg, R. L. (2017). Limb position drift results from misalignment of proprioceptive and visual maps. Neuroscience, 346, 382–394. https://doi.org/10.1016/j.neuroscience.2017.01.040

Penhune, V. B. (2011). Sensitive periods in human development: Evidence from musical training. In Cortex (Vol. 47, Issue 9, pp. 1126–1137). Elsevier. https://doi.org/10.1016/j.cortex.2011.05.010

Philip, B. A., Buckon, C., Sienko, S., Aiona, M., Ross, S., & Frey, S. H. (2015). Maturation and experience in action representation: Bilateral deficits in unilateral congenital amelia. Neuropsychologia, 75, 420–430. https://doi.org/10.1016/j.neuropsychologia.2015.05.018

Philip, B. A., & Frey, S. H. (2011). Preserved grip selection planning in chronic unilateral upper extremity amputees. Exp Brain Res, 214, 437–452. https://doi.org/10.1007/s00221-011-2842-5

Przybyla, A., Coelho, C. J., Akpinar, S., Kirazci, S., & Sainburg, R. L. (2013). Sensorimotor performance asymmetries predict hand selection. Neuroscience, 228, 349–360. https://doi.org/10.1016/j.neuroscience.2012.10.046

Rouder, J. N., Engelhardt, C. R., McCabe, S., & Morey, R. D. (2016). Model comparison in ANOVA. Psychonomic Bulletin and Review, 23(6), 1779–1786. https://doi.org/10.3758/s13423-016-1026-5

Rouder, J. N., Morey, R. D., Speckman, P. L., & Province, J. M. (2012). Default Bayes factors for ANOVA designs. Journal of Mathematical Psychology, 56(5), 356–374. https://doi.org/10.1016/j.jmp.2012.08.001

Rouder, J. N., Speckman, P. L., Sun, D., Morey, R. D., & Iverson, G. (2009). Bayesian t tests for accepting and rejecting the null hypothesis. In Psychonomic Bulletin and Review (Vol. 16, Issue 2, pp. 225–237). Springer. https://doi.org/10.3758/PBR.16.2.225

Rubenstein, J. L. R., Anderson, S., Shi, L., Miyashita-Lin, E., Bulfone, A., & Hevner, R. (1999). Genetic control of cortical regionalization and connectivity. Cerebral Cortex, 9(6), 524–532. https://doi.org/10.1093/cercor/9.6.524

Sainburg, R. L., Lateiner, J. E., Latash, M. L., & Bagesteiro, L. B. (2003). Effects of altering initial position on movement direction and extent. Journal of Neurophysiology, 89(1), 401–415. https://doi.org/10.1152/jn.00243.2002

Sarlegna, F. R., & Sainburg, R. L. (2009). The roles of vision and proprioception in the planning of reaching movements. Advances in Experimental Medicine and Biology, 629, 317–335. https://doi.org/10.1007/978-0-387-77064-2_16

Schneiberg, S., Sveistrup, H., Mcfadyen, B., Mckinley, P., & Levin, M. F. (2002). The development of coordination for reach-to-grasp movements in children. Exp Brain Res, 146, 142–154. https://doi.org/10.1007/s00221-002-1156-z

Scott, S. H. (2004). Optimal feedback control and the neural basis of volitional motor control. Nature Reviews Neuroscience, 5(7), 532–546. https://doi.org/10.1038/nrn1427

Simon-Martinez, C., Lopes Dos Santos, G., Jaspers, E., Vanderschueren, R., Mailleux, L., Klingels, K., Ortibus, E., Desloovere, K., & Feys, H. (2018). Age-related changes in upper limb motion during typical development. https://doi.org/10.1371/journal.pone.0198524

Stankevicius, A., Wallwork, S. B., Summers, S. J., Hordacre, B., & Stanton, T. R. (2020). Prevalence and incidence of phantom limb pain, phantom limb sensations and telescoping in amputees: A systematic rapid review. European Journal of Pain, ejp.1657. https://doi.org/10.1002/ejp.1657

Sur, M., & Rubenstein, J. L. R. (2005). Patterning and plasticity of the cerebral cortex. In Science (Vol. 310, Issue 5749, pp. 805–810). American Association for the Advancement of Science. https://doi.org/10.1126/science.1112070

Sveistrup, H., Schneiberg, S., McKinley, P. A., McFadyen, B. J., & Levin, M. F. (2008). Head, arm and trunk coordination during reaching in children. Experimental Brain Research, 188(2), 237–247. https://doi.org/10.1007/s00221-008-1357-1

Uswatte, G., Taub, E., Morris, D., Light, K., & Thompson, P. A. (2006). The Motor Activity Log-28: assessing daily use of the hemiparetic arm after stroke. Neurology, 67(7), 1189–1194. https://doi.org/10.1212/01.wnl.0000238164.90657.c2

Van Beers, R. J., Haggard, P., & Wolpert, D. M. (2004). The Role of Execution Noise in Movement Variability. Journal of Neurophysiology, 91(2), 1050–1063. https://doi.org/10.1152/jn.00652.2003

Van Beers, R. J., Sittig, A. C., & Denier Van Der Gon, J.J. (1999). Integration of proprioceptive and visual position-information: An experimentally supported model. Journal of Neurophysiology, 81(3), 1355–1364. https://doi.org/10.1152/jn.1999.81.3.1355

Van Den Heiligenberg, F. M. Z., Orlov, T., MacDonald, S. N., Duff, E. P., Henderson Slater, D., Beckmann, C. F., Johansen-Berg, H., Culham, J. C., & Makin, T. R. (2018). Artificial limb representation in amputees. Brain, 141(5), 1422–1433. https://doi.org/10.1093/brain/awy054

van den Heiligenberg, F. M. Z., Yeung, N., Brugger, P., Culham, J. C., & Makin, T. R. (2017). Adaptable Categorization of Hands and Tools in Prosthesis Users. Psychological Science, 28(3), 395– 398. https://doi.org/10.1177/0956797616685869

Vannuscorps, G., & Caramazza, A. (2015). Typical biomechanical bias in the perception of congenitally absent hands. Cortex, 67, 147–150. https://doi.org/10.1016/j.cortex.2015.02.015

Vannuscorps, G., & Caramazza, A. (2016). Typical action perception and interpretation without motor simulation. Proceedings of the National Academy of Sciences of the United States of America, 113(1), 86–91. https://doi.org/10.1073/pnas.1516978112

von Hofsten, C. (1980). Predictive reaching for moving objects by human infants. Journal of Experimental Child Psychology, 30(3), 369–382. https://doi.org/10.1016/0022-0965(80)90043-0

Walton, K. D., Lieberman, D., Llinás, A., Begin, M., & Llinás, R. R. (1992). Identification of a critical period for motor development in neonatal rats. Neuroscience, 51(4), 763–767. https://doi.org/10.1016/0306-4522(92)90517-6

Wesselink, D. B., Heiligenberg, F.M. Van Den Ejaz, N., Dempsey-Jones, H., Cardinali, L., Tarall-Jozwiak, A., Diedrichsen, J., & Makin, T. R. (2019). Obtaining and maintaining cortical hand representation as evidenced from acquired and congenital handlessness. ELife, 8(e37227), 1–19. https://doi.org/10.7554/eLife.37227

Wolpert, D. M. (1997). Computational approaches to motor control. Trends in Cognitive Sciences, 1(6), 209–216.

Wolpert, D. M., Diedrichsen, J., & Flanagan, J. R. (2011). Principles of sensorimotor learning. Nature Reviews Neuroscience, 12(12), 739–751. https://doi.org/10.1038/nrn3112

Wolpert, D. M., Ghahramani, Z., & Jordan, M. I. (1995). An Internal Model for Sensorimotor Integration. Science, 269(29), 1880–1882. http://links.jstor.org/sici?sici=0036-8075%2819950929%293%3A269%3A5232%3C1880%3AAIMFSI%3E2.0.CO%3B2-Q

Zeiler, S. R., & Krakauer, J. W. (2013). The interaction between training and plasticity in the poststroke brain. Current Opinion in Neurology, 26(6), 609–616. https://doi.org/10.1097/WCO.0000000000000025

Zoia, S., Bulgheroni, M., Acus, A., & Skabar, A. (2007). Accessible gaming View project Rewire-Rehabilitative Wayout In Responsive home Environments View project. https://doi.org/10.1007/s00221-006-0607-3

